# Epigenetic control of phenotypic plasticity in a filamentous fungus *Neurospora crassa*

**DOI:** 10.1101/049726

**Authors:** Ilkka Kronholm, Hanna Johannesson, Tarmo Ketola

**Affiliations:** Centre of Excellence in Biological Interactions, Department of Biological and Environmental Sciences, University of Jyväskylä, FI-40014 Jyväskylä, Finland; Department of Organismal Biology, University of Uppsala, 752 36 Uppsala, Sweden

**Keywords:** Reaction norm, DNA methylation, Histone methylation, Histone deacetylation, RNA interference, fungi

## Abstract

Phenotypic plasticity is the ability of a genotype to produce different phenotypes under different environmental or developmental conditions. Phenotypic plasticity is an ubiquitous feature of living organisms, and is typically based on variable patterns of gene expression. However, the mechanisms by which gene expression is influenced and regulated during plastic responses are poorly understood in most organisms. While modifications to DNA and histone proteins have been implicated as likely candidates for generating and regulating phenotypic plasticity, specific details of each modification and its mode of operation have remained largely unknown. In this study, we investigated how epigenetic mechanisms affect phenotypic plasticity in the filamentous fungus *Neurospora crassa*. By measuring reaction norms of strains that are deficient in one of several key physiological processes we show that epigenetic mechanisms play a role in homeostasis and phenotypic plasticity of the fungus across a range of controlled environments. Effects on plasticity are specific to an environment and mechanism, indicating that epigenetic regulation is context dependent and is not governed by general plasticity genes. In our experiments with *Neurospora*, histone methylation and the RNA interference pathway had the greatest influence on phenotypic plasticity, while lack of DNA methylation had the least.

## Introduction

Natural environments are in a constant state of change. Organisms must cope with different environments and their dynamism by adjusting their development, behavior and reproduction while always maintaining a physiological homeostasis. In most cases, phenotypic plasticity is based on adjusting the patterns of gene expression (Nicotra *et al*., 2010). Phenotypic plasticity plays an important role in buffering fitness across a range of environments, and can help evolutionary adaptation to extreme environments (Lande, 2009; Chevin *et al*., 2010; Draghi and Whitlock, 2012). Specifically, plasticity can facilitate adaptation by increasing population size in the novel environment (Chevin *et al*., 2010) and by generating phenotypic variation that can be selected on if it is heritable (PÁl, 1998). Heritable phenotypic variation could potentially be achieved with different mechanisms, one of them being plasticity mediated by epigenetic mechanims. Interestingly, evolutionary models incorporating heritable genetic and epigenetic systems suggest that the latter play a significant role in adaptation (Day and Bonduriansky, 2011; Klironomos *et al*., 2013; Kronholm and Collins, 2015).

Transcriptomic studies have shown that environment has a large effect on gene expression profiles (Gibson, 2008; Nicotra *et al*., 2010; Alvarez *et al*., 2015). The classical view is that those patterns are regulated by transcriptional activator and repressor proteins (Davidson, 2006). However, there is increasing interest in the influence of epigenetic changes (e.g., DNA methylation and histone modifications) on gene expression (de la Paz Sanchez *et al*., 2015). Furthermore, epigenetic mechanisms are involved in phenotypic plasticity (Slepecky and Starmer, 2009; Boss-Dorf *et al*., 2010; Herrera *et al*., 2012; Baerwald *et al*., 2015) and transgenerational inheritance(Verhoeven *et al*., 2010; Verhoeven and van Gurp, 2012; Luna and Ton, 2012; Rasmann *et al*., 2012; Ou *et al*., 2012; Öst *et al*., 2014; Siklenka *et al*., 2015). While experiments that have artificially induced variation in DNA methylation have shown how this factor can contribute significantly to phenotypic variation (Cortijo *et al*., 2014) and plasticity (Kooke *et al*., 2015), the relative importance of other epigenetic modifications remains unclear - even the extent to which their effects are environmentally dependent.

To explore the extent to which various epigenetic mechanisms are involved in phenotypic plasticity, we used an experimental set of 25 different mutant strains of the filamentous fungus *Neurospora crassa*, each of which was deficient in a particular chromatin modification, affecting: DNA methylation (3 mutants), histone methylation (5), histone deacetylation (8), histone acetylation (2), histone demethylation (2), and RNA interference (5). We selected mutants that either had been previously characterized and were known to affect different epigenetic modifications or based on their homology to genes known to modify chromatin in other organisms.

We measured the reaction norms of each strain with respect to four different environmental variables: temperature, osmotic stress (NaCl), sucrose concentration, and pH to investigate the potential effects of each epigenetic mechanism on phenotypic plasticity across a range of environments. A reaction norm is visualized by plotting the measurable performance (e.g., mycelial growth rate) of an organism scored at different values of an environmental parameter. Reaction norms can be described by their shape and elevation; shape refers to variation in phenotype that contributes to genotype by environment interaction and elevation means variation in phenotype that contributes to the genotypic effect only. If epigenetic modifications play a role in phenotypic plasticity, we expect to see differences reaction norms shape for the mutant strain and wild type. Differences in reaction norm elevation indicate that the epigenetic modification is required for normal cellular function rather than a physiologically-plastic response.

We show that epigenetic mechanisms are involved in plastic responses of *N. crassa*. These responses involve a specific epigenetic modification in a particular environment, e.g., histone modifications are important in the response to temperature and pH, and the RNA interference pathway also has notable effects. In contrast, lacking the ability to carry out DNA methylation had little effect on strain performance in any of the trial environments.

## Materials and methods

### Neurospora strains

We used 25 different strains from the *N. crassa* knockout collection (Colot *et al*., 2006) to investigate the role of epigenetic mechanisms in phenotypic plasticity. The mutants were generated by replacing the entire open reading frame of the target gene with a *hph* cassette, which confers resistance to the antibiotic Hygromycin B. Strains were obtained from the Fungal Genetics Stock Center (FGSC) (Mccluskey *et al*., 2010); table S1 shows the strains used in this study and table 1 shows the genotypes of those used in the experiments. We included strains that were viable in the homokaryotic state and for which we could confirm the gene deletion. Strain FGSC # 4200 was used as a wild type control for the reaction norm measurements. We grouped the strains into five categories based on the epigenetic mechanism for which they are deficient: DNA methylation, histone methylation, histone deacetylation, RNA interference, and “other” which included two putative histone demethylases and two histone acetyl transferases (Table S1).

**Table 1:**
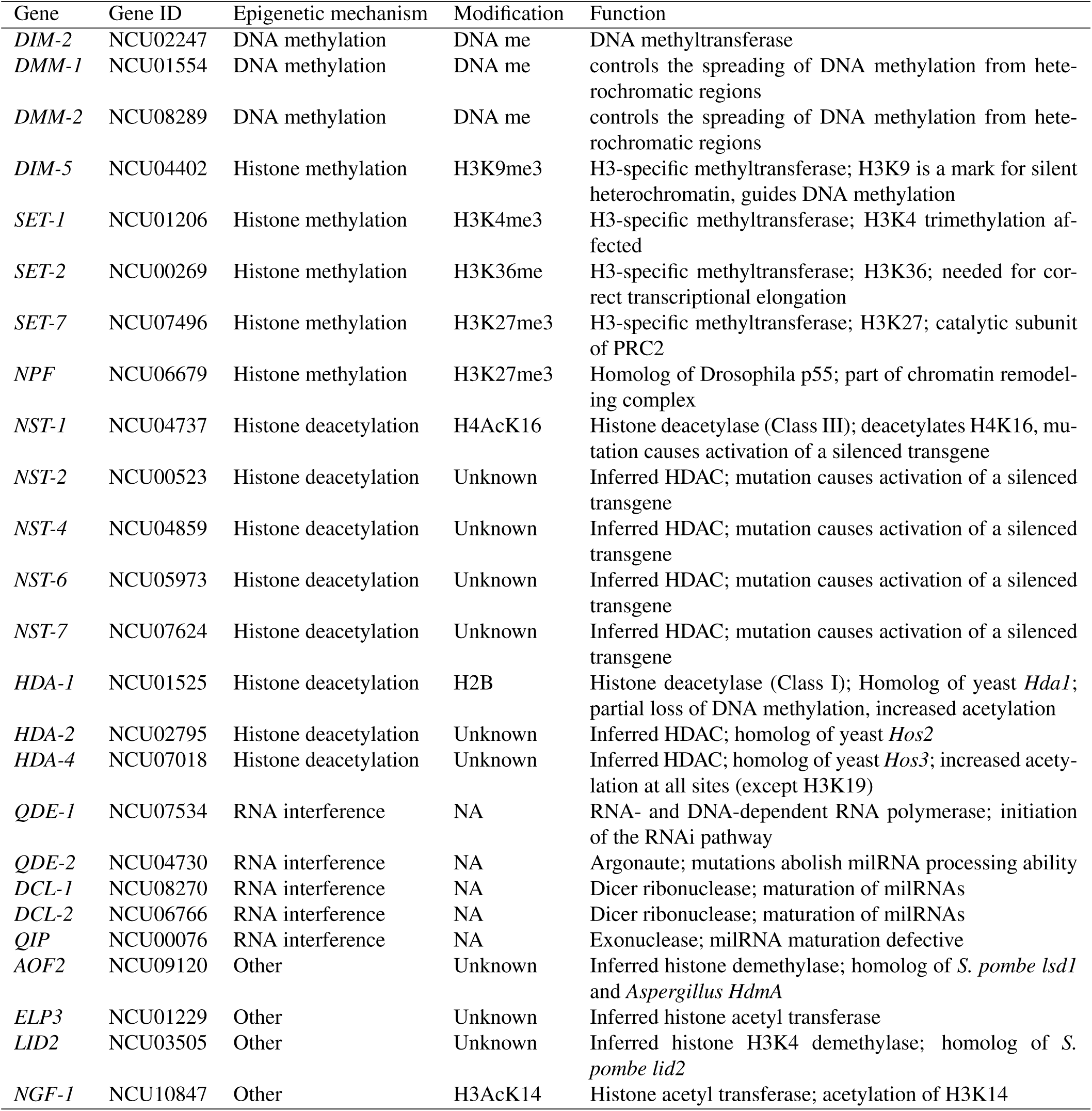
Mutant strains used in this study. Gene ID is based on the *N. crassa* genome assembly NC12. Epigenetic mechanism is the classification for the strains studied here. Modification is the chromatin modification affected by the mutation and function describes what is known about the biochemical activity of the protein. NA = not applicable, HDAC = histone deacetylase, milRNA = micro RNA-like RNA

### Genotyping

We confirmed that the mutant strains indeed had deletions and verified their mating types by PCR. We relied on a rapid method of DNA extraction from conidia, or asexual spores (Henderson *et al*., 2005). We grew a strain in an agar slant for 3–5 days until orange-colored spores were visible. Spores were then collected and suspended in water containing 0.01 % Tween-80. Conidial boiling buffer was prepared by combining 100 parts of 50 mmol/l Tris pH 8 and 2 parts of 0.5 mol/l EDTA pH 8.5. We then distributed 10 μl of conidial boiling buffer across a 96-well PCR plate and combined 40 μl of conidial suspension to each well. The loaded plate was boiled for 10 minutes at +98 °C in a thermal cycler, and 2 μl of the resulting suspension was used as template in subsequent PCR reactions.

Strain mating type was determined by PCR in a single 10 μl reaction containing four primers, i.e., two pairs. These two primer pairs (Table S2) were designed using the *N. crassa* genome sequence to amplify a 200-bp fragment from the *mat a* locus and a 400-bp fragment from the *mat A* locus. PCR products were resolved by electrophoresis on an 2–3 % agarose gel. To confirm that our mutant strains indeed had deletions, we designed primers for each of the target genes that amplified an approximately 500-bp fragment and checked the presence of the deleted genes by PCR. We set up PCR reactions with the high-fidelity Phusion DNA polymerase (Thermo Scientific) according to manufacturer’s instructions: the final PCR reactions contained 1x Phusion HF buffer, 0.2 mmol/l each dNTP, 0.5 μmol/l each primer, and 0.2 or 0.4 units of Phusion DNA polymerase for 10 or 20 μl reactions, respectively. All PCR reactions were run with the following thermal profile: initial denaturation of +98 °C for 30 s, then 35 cycles of +98 °C for 5 s, variable annealing for 10 s, extension at +72 °C for 20 s, and a final extension at +72 °C for 1 min. Primer sequences and their annealing temperatures are given in table S2.

### Growth measurements

In general, standard laboratory protocols for *Neurospora* (Davis and de Serres, 1970) were followed. We grew *N. crassa* on Vogel’s growth medium (Metzenberg, 2003) with 1.5 % agar appropriately supplemented for the different environments listed below. To measure growth rate of the strains, we used a race tube method of Ryan *et al*. (1943) with tubes prepared following White and Woodward (1995). Briefly, we filled 25 ml plastic serological pipettes (Sarsted) with 10 ml of molten agar and placed them horizontally so that the agar solidified at the bottom of the pipettes. The tip of the pipette was snapped off, the end inoculated with conidia of a given strain, and sealed with parafilm. Growth of the mycelial front in the tube was measured by marking its position twice a day (every 8th and 16th hour). Growth was followed typically for a period of 104 hours but up to 152 hours in the osmotic stress environment.

To estimate growth rates we used a simple linear regression of time against the distance the mycelial front had grown in a race tube. Growth data were collected from the first mark after inoculation, i.e., growth immediately following inoculation, until the mycelial front was first visible was not included. This effectively corrects for possible differences in initial growth rate due to inoculum size. We extracted the slope of the regression line for each growth assay to obtain the mycelial growth rate as mm/h. Growth rates were used as a dependent variable in subsequent analyses.

### Reaction norms

As a measure of phenotypic plasticity, we measured reaction norms of the different strains with respect to four different environmental parameters: temperature, osmotic stress (NaCl), sucrose concentration, and pH. We used six different settings within each parameter, 26 different genotypes with five replicate growth rate measurements in each treatment combination, yielding 3120 growth rate measurements in total. The different parameter settings were: +15, +20, +25, +30, +35, +40 °C for the temperature; 0, 0.2, 0.4, 0.8, 1.2, 1.6 mol l^™1^ of NaCl added for osmotic stress; 0.015, 0.15, 1.5, 5, 15, 30 % (weight/volume) of sucrose added for sucrose concentration; or pH adjusted to 4.0, 5.0, 5.8, 7.0, 8.0 and 9.0. Except for temperature, which was controlled by the growth chamber, the standard growth medium was manipulated by either adding NaCl, varying the sucrose level or adjusting pH with either HCl or NaOH. We did not control for any changes in nutrient availability in the medium that may result from pH changes, and allowed the environmental changes to be complex.

Normal growth conditions were +25 °C, 0 mol l^™1^ NaCl, 1.5 % sucrose, and pH 5.8, and independent measurements were collected during all experiments under these “control” conditions. The reaction norm experiment was performed in growth chambers and replicate measurements were blocked in time by replicates, such that each strain and environmental setting ran simultaneously and the growth tubes were randomized in the growth chamber. As the temperature treatment had to be applied to the entire growth chamber, we used two growth chambers of the same model (Lab companion ILP-02/12; Jeio Tech, South Korea) where we always switched the identity of the growth chamber between replicate measurements. This allowed us to check for any possible effect of either growth chamber independent of temperature.

### Backcrossing and validation

While the genetic background of mutant strains in the knockout collection and the control should be nearly isogenic (Colot *et al*., 2006), there remains the possibility that some genetic background effects could be present. To rule these and any other subtle phenotypic effects out, we performed a validation experiment with strains where we had backcrossed the mutant strain five times into FGSC # 2489 using standard crossing techniques (Davis and de Serres, 1970). The genetic background of strains 2489 and 4200 is nearly isogenic, except for the mating type locus (Mylyk *et al*., 1974). And as the mutant strains share their background with 2489 / 4200 (Colot *et al*., 2006) the mutant strains and the controls share more of their genome than would be expected from five backcrosses.

The sexual cycle can be induced by growing *N. crassa* in a low-nitrogen medium. The fungus has two different mating types: *mat A* and *mat a*. When the two types meet they fertilize each other and undergo meiosis. We used crossing medium (Davis and de Serres, 1970) containing 0.2 % sucrose but instead of agar we used 5 ml of liquid medium in a large 20 * 150 mm test tube with a 40 * 90 mm piece of vertically-folded filter paper that wicks the medium by capillary action. The filter paper was inoculated with conidia from the opposite mating types. We isolated ascospores on plates with sorbose to induce colonial growth and 200 μg/ml of Hygromycin B, where appropriate, to select for mutant-strain progeny. We checked mating type of the progeny and confirmed the gene deletions by PCR in each round of backcrossing as above. As 2489 is mat A we selected mat a progeny until a final backcross generation was recovered containing mutant genotypes of both mating types.

Based on the results of the reaction norm experiment, we selected strains *dim-2*, *dmm-2*, *hda-1*, *qde-2*, *qip*, *aof2*, *lid2*, and *set-7* for validation experiments with the backcrossed strains. For temperature measurements we measured *dim-2*, *qip*, *set-7*, and *aof2* in +40 °C, *qde-2*, *lid2*, and *aof2* in +35 and +30 °C. For the pH environment we measured *dmm-2* at pH 4, and *dmm-2* and *qde-2* at pH 9. In osmotic stress we only measured *hda-1* at 0.8 mol l^™1^ NaCl and for sucrose concentration we measured *dim-2*, *qde-2*, and *set-7* at 30 % sucrose, and *qde-2* at 0.015 % sucrose. Growth rate was measured as in the reaction norm experiment, and assays were replicated 12 times in the validation experiment, each including strain 2489 as a control.

### Data analysis

We used an ANOVA to investigate whether different epigenetic mechanisms have different effects and whether these are specific to particular environments. Because the reaction norms were non-linear, we encoded the different parameter settings as factors. This allowed us to analyze all of the data together despite differences in reaction norm shape. We fitted a mixed model using the ‘lmer’ function in R (R Core Team, 2013) with tests performed using the ‘lmerTest’ package (Kuznetsova *et al*., 2015). This package implements F-tests using type III sums of squares with Satterwhaite correction for degrees of freedom. Type III sums of squares were used to interpret results according to the elevation and shape of the reaction norms, following the phenotypic plasticity literature. The model was

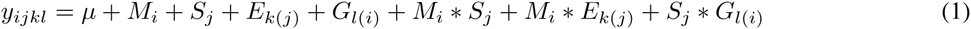

where *μ* is the intercept, *M*_*i*_ is the *i*th epigenetic mechanism, *S*_*j*_ is the *j*th environmental parameter (temperature, salt, sucrose or pH stress), *E*_*k*(*j*)_ is the *k*th parameter setting nested within parameter *j* and *G*_*l*(*i*_) is the *l*th genotype nested within epigenetic mechanism *i*. Epigenetic mechanism, environmental parameter, and parameter setting were fitted as fixed factors, while genotype was fitted as a random factor. We subsequently analyzed each of the different environmental parameters separately with the model

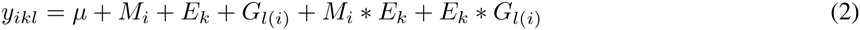

with terms the same as above.

### Estimating reaction norm optima

To characterize reaction norm optima for the different strains we fitted natural splines, i.e. functions built piecemeal from polynomial functions (Venables and Ripley, 2002), to each of the genotypes in each of the environmental parameters, as implemented in the ‘splines’ R package. For this and subsequent analyses we encoded the different parameter settings as continuous variables, we used splines because some of the strains showed reaction norm shapes that made fitting the same regression model to each of the genotypes inappropriate. The drawback of using splines is that we cannot estimate the critical thresholds when growth rate approaches zero, as natural splines do not allow extrapolation outside of the data range.

### Bayesian estimation of differences between the control and mutant strains

To test for differences between the control and mutant strains in specific environments, we used a Bayesian model analogous to a one-way ANOVA. The model specification followed Gelman (2006) and Kruschke (2011):

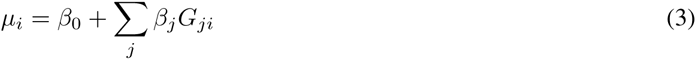

where *β*_0_ is the grand mean, *β*_*j*_ is the effect of the *j*th genotype *G*. For the analysis we standardized the data such that *β*_0_ = 0 and ∑_*j*=1_ *β*_*j*_ = 0. Observations are assumed to be distributed normally around *μ*_*i*_, *y*_*i*_ ~ *N*(*μ*_*i*_, *τ*_*j*_), where *τ*_*j*_ is a precision of the normal distribution for the *j*th genotype. We used a hierarchical prior for estimating *β*_*j*_ for each setting of *G* following Gelman (2006), *β*_*j*_ ~ *N*(0, *τ*_*β*_), where 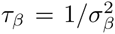 and we used a folded t-distribution as a prior for *σ*_*β*_. Since we are allowing each setting of *G* to have its own variance, *τ*_*j*_ is distributed as *τ*_*j*_ ~ Γ(*s*_*G*_, *r*_*G*_), where *s*_*G*_ = *m*^2^/*d*^2^, *r*_*G*_ = *m*/*d*^2^ and *m* ~ Γ(*s*_*m*_, *r*_*m*_) and *d* ~ Γ(*s*_*d*_, *r*_*d*_). The benefit of using Bayesian analysis here instead of classical ANOVA is that we allow each genotype to have its own variance and that there is no special adjustment needed for multiple comparisons as the hierarchical model takes this into account via shrinkage (Gelman, 2006; Gelman *et al*., 2012). The model was implemented following Kruschke (2011) with the JAGS program (Plummer, 2003) using the R package ‘rjags’ (Plummer, 2014). For the MCMC sampling we used five chains with a burn-in of 10000 iterations and sampled 10000 iterations while thinning by 750 to remove auto-correlation between samples. We checked convergence of the MCMC simulation through graphical diagnostics.

We calculated contrasts between the control and mutant strains using the posterior distributions for *β*_*j*_ as *β*_*mutant*_ – *β*_*control*_, we also re-scaled the *β*_*j*_ values to their original scale. If a strain is growing slower than the control, their difference will be negative. We considered growth rates of the control and the mutant strains to be significantly different if the 95 % highest posterior density (HPD) interval for the difference did not include zero.

### Data availability

Strains are available upon request. File S1 contains all phenotypic data.

## Results

### Growth rates

We found that growth of *N. crassa* in our race tubes to be linear: the 95 % quantiles for *R*^2^ values of a linear fit across all measurements were 0.952 – 0.999 with a median of 0.998. The only genotypes exhibiting deviations from a strict linear pattern were *dim-5*, *ngf-1*, and *npf*, the latter showing the lowest *R*^2^ in the whole dataset (0.622). This deviation from linear growth was observed particularly in environments where growth was very slow. Because these cases represent only a small portion of the entire dataset, we also used a linear model for the growth of these genotypes. We found that in the control environment our control genotype 4200 grew at a rate of 3.29 ±0.24 (95 % CI) mm/h, in line with previous reports (Ryan *et al*., 1943).

In some tubes where no growth had occurred during the growth assay we could observe growing mycelium after an extended amount of time, (e.g. *npf* in high osmotic stress and *dim-5* at low temperature). We assigned a growth rate of 0 to those measurements.

### Reaction norm experiment

In the reaction norm experiment, data was missing for 19 out of 3120 measurements. Most of these were probably due to failed inoculations, as in many cases we were able to distinguish between missing data and no growth in a particular trial. Reaction norms were visualized by plotting growth rate against the different environmental parameters (Figure 1). Visual inspection revealed that nearly all reaction norms were non-linear, even in the osmotic stress reaction norms there was some indication of curvature. Even if in the osmotic stress environment the optimum is at zero and growth rate decreases as salt concentration increases (Figure 1). First, we performed an ANOVA to investigate whether the different epigenetic mechanisms have different effects on phenotypic plasticity (Table 2). We did not observe a significant main effect of epigenetic mechanism type, but we did find a significant interaction between epigenetic mechanism and parameter setting nested within stress type, *F*_80,400.17_ = 1.397, *p* = 0.021 (Table 2). This result indicates that different epigenetic mechanisms have different effects on different environmental parameters. Thus, we subsequently analysed the data by stress type.

**Figure 1:**
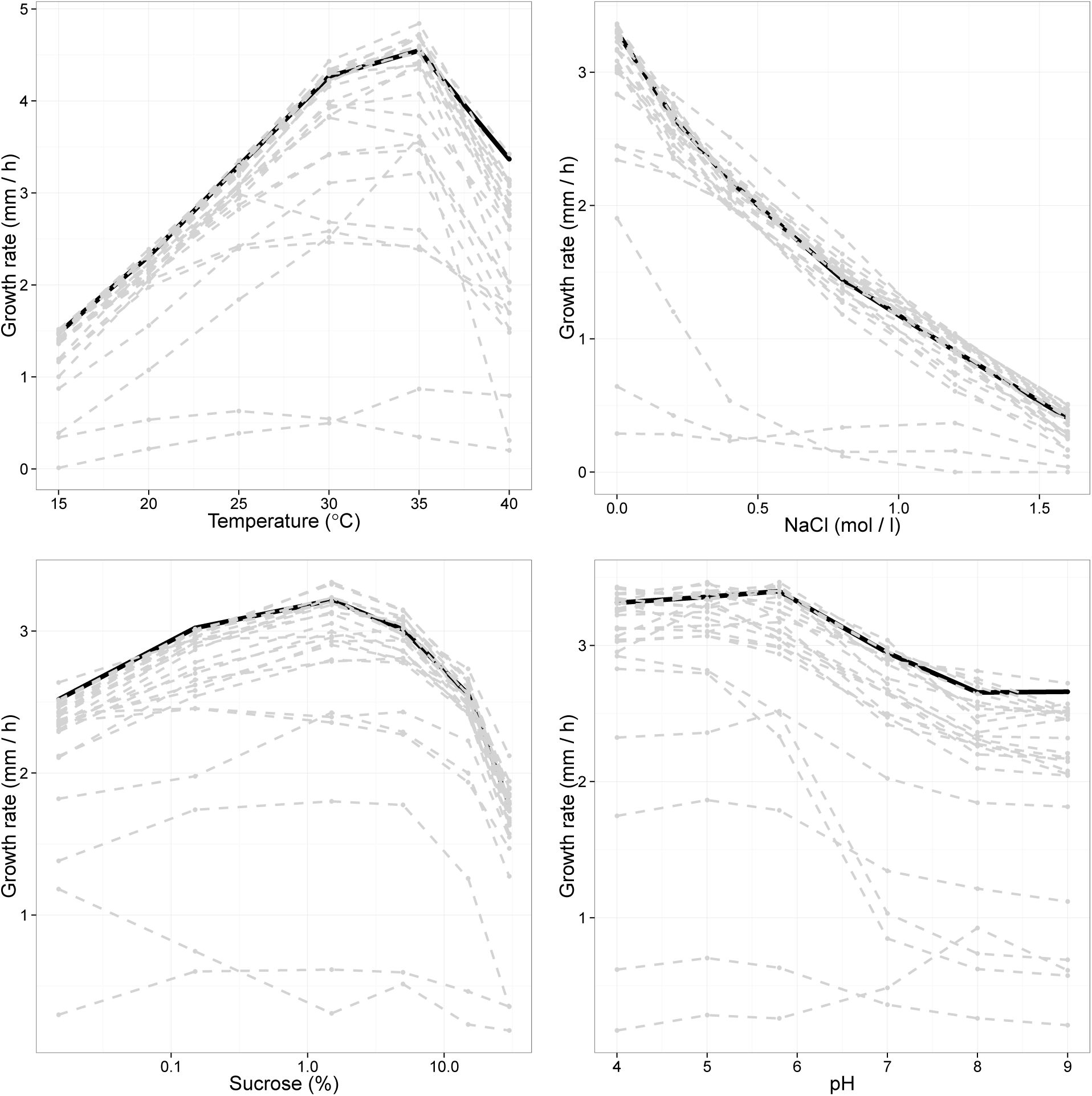
Overview of reaction norms according to each environmental parameter: temperature; osmotic stress; sucrose concentration; and pH (from top left to bottom right). Horizontal axis shows the parameter setting and vertical axis the growth rate. Black reaction norms are the control and dashed gray lines are the different mutant strains. See supplementary material for detailed pictures of the different epigenetic mechanisms.

**Table 2:** ANOVA table of overall differences in reaction norms for all data. Fixed effects were tested with F-tests with Satterthwaite approximation for degrees of freedom and random effects were tested with *χ*^2^-test. For fixed effects: Df = numerator degrees of freedom, Df2 = denominator degrees of freedom.

| Fixed effects | Df, Df2 | F value | p-value |
| --- | --- | --- | --- |
| Mechanism | 4, 20.02 | 2.189 | 0.107 |
| Environmental parameter | 3, 60.03 | 126.668 | <2E-16 |
| Environmental setting (within E. param.) | 20, 400.37 | 149.157 | <2E-16 |
| Mechanism * Environmental parameter | 12, 60.02 | 1.22 | 0.291 |
| Mechanism * Environmental setting (within E. param.) | 80, 400.17 | 1.397 | 0.021 |
| Random effects | $\chi^2$ | Df | p-value |
| Genotype | 110.6 | 1 | <2E-16 |
| Environmental parameter * Genotype | 36.4 | 1 | <2E-9 |
| Environmental setting (within E. param.) * Genotype | 1852.1 | 1 | <2E-16 |

When analyzing the environmental parameters separately, we did not observe a significant main effect of epigenetic mechanism in any of them and the interaction between mechanism and parameter setting was only significant in the pH trial *F*_20,100.061_ = 2.271, *p* = 0.004. However, the main effect of genotype and the interaction between genotype and parameter setting is significant for each parameter (Table 3). These results suggest that epigenetic mechanisms in general contribute to phenotypic plasticity. Among the genotypes there were differences in both elevation and shape of the reaction norms. We did not see a general effect of particular epigenetic modifications type, such as all mutant strains with nonfunctional histone methylation have a characteristic reaction norm. On the contrary, we noticed that the reaction norms were specific to a given mutant strain in a particular environment. We also performed the previous analyses excluding genotypes *ngf-1* and *dim-5* as these two genotypes grew much slower in general than rest, but this did not change any of our conclusions. For the temperature trial, we also investigated whether the two growth chambers used had any different effects on growth. We included growth chamber identity as a fixed factor in the mixed model for the temperature trial, but this term was not significant *F*_1,593.15_ = 0.545, p = 0.461 and growth chamber term was dropped from the final model.

**Table 3:**
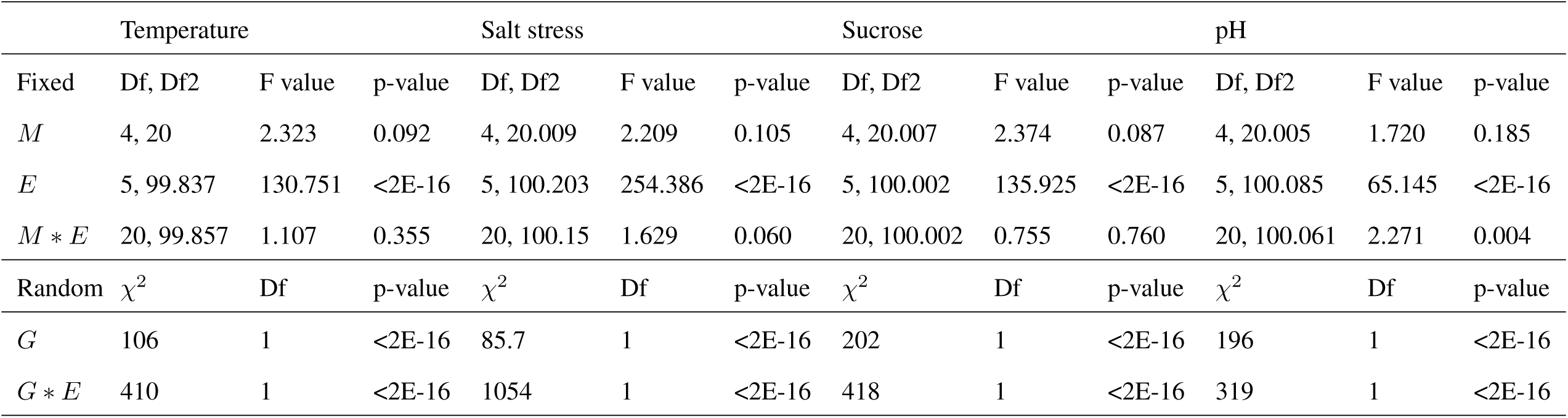
ANOVA table of overall differences in reaction norms separately for each environmental parameter. *M* = Mechanism, *E* = Environmental setting, *G* = Genotype, Fixed effects were tested with F-tests with Satterthwaite approximation for degrees of freedom and random effects were tested with *χ*^2^-test. For fixed effects: Df = numerator degrees of freedom, Df2 = denominator degrees of freedom.

We also observed lhat there were changes in reaction norm optima among the mutant strains as calculated from the natural spline fits. Plotting the distribution for the optimal environment of each genotype shows that osmotic stress is the only parameter where the optimum remains constant at 0 mol/1 NaCl for all genotypes (Figure 2), except for *dim-5* where the reaction norm is rather flat (Figure S2). In the other environments some genotypes have different environmental optimum than the control (Figure 2).

**Figure 2:**
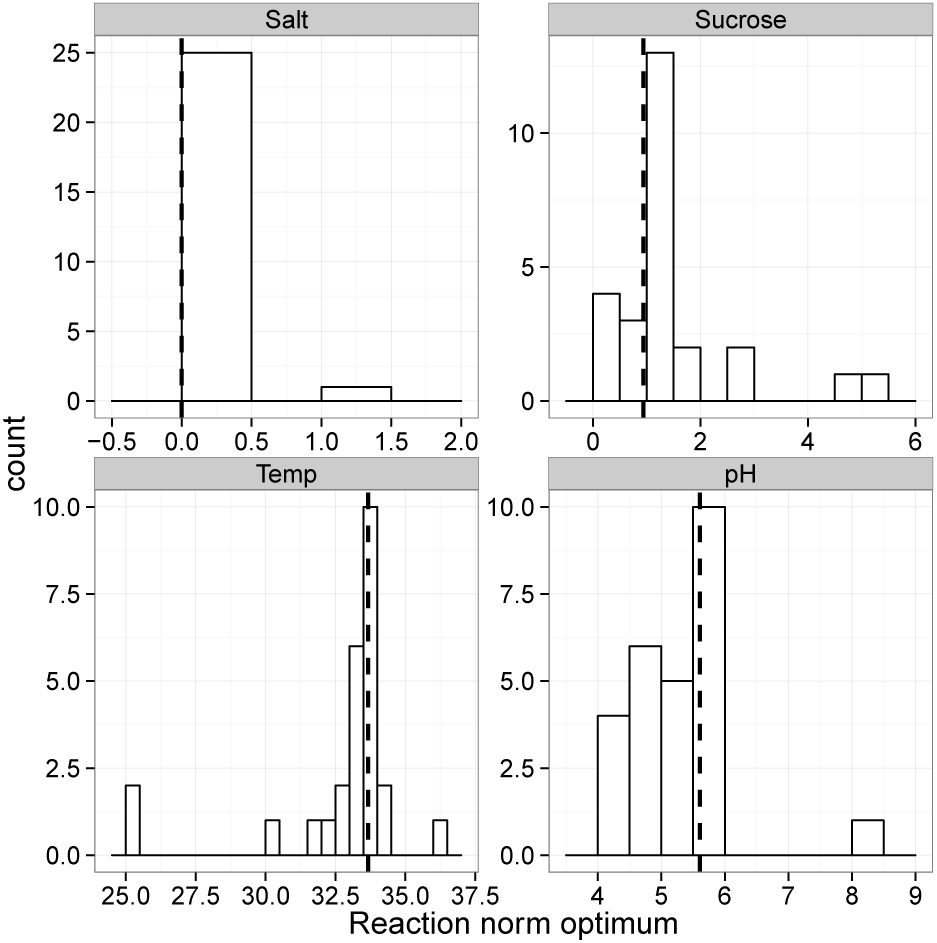
Distributions of the optimal settings for the mutant strains for the four environmental parameters. An optimal environment was estimated from a natural spline fit to the reaction norm data for each strain. Dashed vertical lines indicate the optimal parameter setting for the control.

Having established that there are statistically-significant differences among the mutant strains in the different environments, we now present the results according to each epigenetic mechanism and by parameter highlighting interesting mutants and their effects.

### Effects of DNA methylation

So far, no apparent phenotype other than the lack of methylation has been detected for the DNA methyltransferase mutant *dim-2*. In our experiment, *dim-2* failed to demonstrate any observable phenotypic effects in the pH, salt, or sucrose trials. In the temperature trial, *dim-2* had no effect up to +35 °C but grew slower than the control at +40 °C; difference to the control –0.57 (–1.14 to – 0.02, 95% HPD) mm/h and we sought to validate this suggestive difference. In the validation experiment we observed that *dim-2* grew at the same rate as the control; difference 0.01 (–0.17 to 0.19, 95% HPD) mm/h. Thus, we conclude that lacking a DNA methylation system has no effect at +40 °C. In the 30 % sucrose setting, we first observed a non-significant tendency of *dim-2* to grow faster than the control (Figure S1), and we confirmed this finding in a validation experiment where we observed a small but significant effect; a difference of 0.07 (0.03 to 0.12) mm/h.

The *dmm-1* and *dmm-2* mutants control the propagation of DNA methylation from heterochromatic regions (Honda *et al*., 2010). We observed that while *dmm-1* had the same optimum sucrose concentration as the control, it grew slower at lower sucrose concentrations (Figure S1); difference to control at 0.015 % sucrose –0.21 (–0.38 to – 0.05) mm/h – a reduction of 8 %. It also grew slower at higher pH and the reaction norm had a lower elevation in the salt stress trial. Generally, *dmm-2* had a slightly lower elevation for all reaction norms (Figure S1). In the pH trial, its reaction norm had a different shape compared to the control and the optimal setting of *dmm-2* was 5.0 compared to 5.8 of the control and the difference in growth rates at pH 4 was –0.36 (–0.61 to – 0.11) mm/h. We validated the growth of *dmm-2* at pH 4 (difference was –0.43 (–0.50 to –0.36) mm/h) and at pH 9 (difference was –0.61 (–0.70 to –0.51) mm/h). Confirming the phenotypic response to different pH for *dmm-2*. Thus, the dmm mutant strains had different phenotypic responses and the results suggest that (possibly silenced) genes adjacent to heterochromatic regions control the response to several environmental parameters.

Overall, our data suggest that DNA methylation does not play a very important role in phenotypic plasticity, but spurious DNA methylation has the potential to affect phenotypic plasticity.

### Effects of histone methylation

Histone methylation has multiple functions depending on which residues are methylated (Rothbart and Strahl, 2014). For the *set-7* mutant, which lacks H3K27me3, we did not observe any growth responses among the environmental parameters and settings we tested (Figure S2). The *npf* mutant also lacks H3K27me3 but also has other functions (Jamieson *et al*., 2013). Generally, *npf* grew much slower than the control; in the sucrose and pH trials the differences were seen in reaction norm elevation rather than shape (Figure S2) but in the temperature and salt stress trials *npf* presents a different shape. In particular, *npf* seems to be sensitive to high temperatures and osmotic stress as its growth rate collapses at +40 °C and in high salt (Figure S2). However, these changes are not related to H3K27me3 as *set-7* does not show them.

The *set-1* mutant lacks H3K4me3 and also shows a lower elevation of its reaction norms (Figure S2) in the sucrose and pH trials: growth of *set-1* slows down more than for the control when pH is changed from 5.8 to pH 4.0 (a difference of 0.19 [–0.05 to 0.43) mm/h), and for the control the difference is only 0.09 (–0.15 to 0.32) mm/h, although this is only marginally significant. This can also be observed when going from 1.5 % sucrose to 0.15 % sucrose; for *set-1* the difference is 0.37 (0.17 to 0.58) mm/h while for the control the difference is only –0.08 (–0.28 to 0.12) mm/h. As H3K4me3 has been implicated in transcriptional activation (Pokholok *et al*., 2005; Raduwan *et al*., 2013), some genes may not activate correctly in these environments.

H3K36me is believed to be required for efficient transcriptional elongation (Morris *et al*., 2005). For the set-2 mutant which lacks H3K36me, we observed a generally lower elevation for reaction norms (Figure S2) but not to the same extent as *set-1*. However, *set-2* has a markedly different response to temperature as its optimal setting from a natural spline fit is at +25.3 °C compared to the +33.7 °C of the control (Figure S2). This suggests that H3K36me is involved in the transcription of genes required for a high temperature response.

The final histone modification we investigated was H3K9me; the mutant *dim-5* that lacks this modification (Tamaru and Selker, 2001; Tamaru *et al*., 2003) grew very poorly in all environments (Figure S2), as noted previously (Tamaru and Selker, 2001). Thus H3K9me plays a central role in essential cellular processes.

Taken together, H3K27 trimethylation has no observable phenotypic effect, H3K4 trimethylation and H3K36 methylation have some effects on elevation of reaction norms but also on their shapes, while H3K9 methylation is needed for normal cellular function.

### Effects of histone deacetylation

For histone deacetylation we used two different classes of mutants: *hda-1*, *hda-2*, and *hda-4* and type III (NAD^+^ dependent) histone deacetylases *nst-1*, *nst-2*, *nst-4*, *nst-6*, and *nst-7*. We observed that for the hda mutants, reaction norm elevations were reduced (Figure S3), except in the salt stress trial where *hda-1* and *hda-2* grew faster than the control (Figure 3). The difference in growth between *hda-1* and the control was –0.19 (0.07 to 0.31), an increase of 13 %. Such an increase could also be observed in the 30 % sucrose environment.

**Figure 3:**
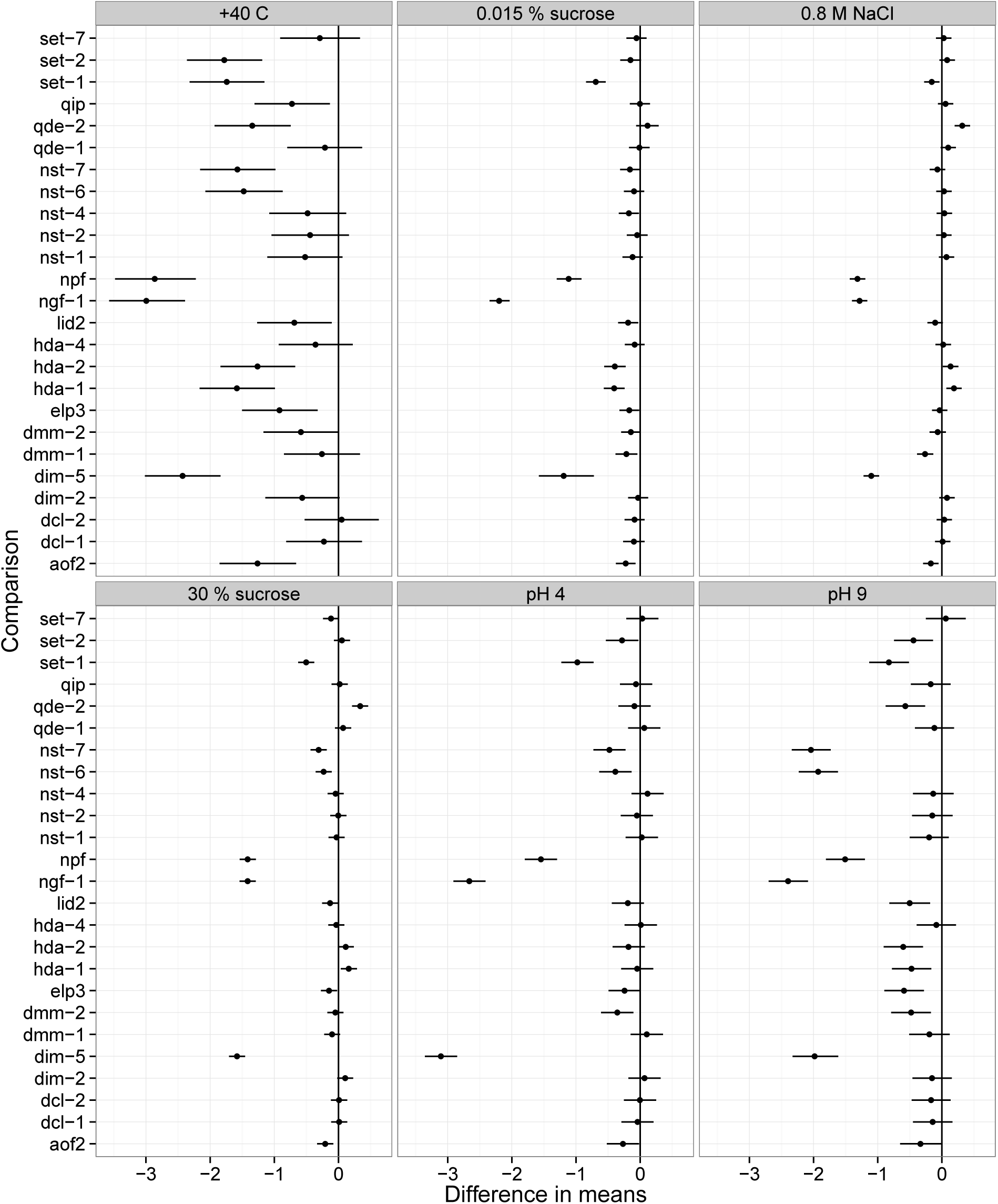
Contrasts for the mutant growth rates in different environments. Vertical axis shows the different mutants and the horizontal axis shows the mutant – control difference in mean growth rates, error bars are 95 % HPD intervals. Vertical lines in the panels show the difference of zero line.

For the *nst* mutants 1, 2, and 4 we did not observe any notable phenotypic effects. The two remaining *nst* mutants (6 and 7) showed large growth effects in nearly all trials. Their effects were particularly noticeable in the pH trial where growth was drastically reduced at high pH (Figure S4, Figure 3) and both of their optimal environments were at lower pH settings than for the control: pH 4.0 for *nst-6* and pH 4.5 for *nst-7* compared to the optimum of pH 5.6 of the control. Also, their reaction norms to sucrose at low concentrations were flat, they did not increase their growth rate in response to rising sucrose concentrations from 0.015 % sucrose as the control does.

Thus, *hda-1* and *hda-2* generally presented a lower reaction norm elevation similar to *nst-6* and *nst-7*, and which also had a different reaction norm shape. Otherwise, histone deacetylation mutants showed no observable responses in terms of their reaction norms and the parameters and settings we tested.

### Effects of RNA interference

For the RNA interference mutants we did not observe any effects of the two Dicer genes, *dcl-1* and *dcl-2*, or the *qde-1* gene (Figure S5). For the *qip* mutant we only saw a phenotype in the +40 °C environment, where the difference to control was –0.72 (–1.31 to – 0.13) mm/h. However, when we attempted to replicate this finding in a validation experiment we did not observe any significant differences between *qip* and the control; the difference was 0.03 (–0.15 to 0.21) mm/h. Thus, we conclude that *qip* has no effect at +40 °C.

However, we observed a different reaction norm shape in all trials with *qde-2* (Figure S5). It grew faster than the control at intermediate salt concentrations (Figure 1), for instance in 0.8 M NaCl its difference in growth rate to the control was 0.32 (0.20 to 0.44) mm/h – an increase of 22 %. An increase in growth rate compared to the control was also observed in the 30 % sucrose environment, a difference of 0.34 (0.21 to 0.46) mm/h. We confirmed this result in a validation experiment where we observed a difference of 0.33 (0.28 to 0.38) mm/h. The growth rate of *qde-2* also had a tendency to increase in the 0.015 % sucrose environment and although this effect was not significant in our first experiment, we observed a significant increase in the validation experiment; a difference of 0.10 (0.05 to 0.16) mm/h. Based on natural spline fit, the *qde-2* mutant strain also had a lower optimal temperature of +31.9 °C (versus + 33.7 °C), and we validated this result by measuring backcrossed *qde-2* at+30 and +35 °C. We observed that the shape of the *qde-2* reaction norm did indeed change and the growth rate of *qde-2* increased by 0.29 (0.18 to 0.41) mm/h compared to the an increase of 0.68 (0.57 to 0.79) mm/h when the temperature increased from +30 to +35 °C. This indicates that microRNA-like molecules that *Neurospora* produces (Lee *et al*., 2010) are involved in the response to high temperatures. For the pH trial, elevation of the *qde-2* reaction norm was generally lower but at pH 9 *qde-2* a shape change was indicated (Figure S5). The growth difference between the control and *qde-2* at pH 9 was validated (difference of –0.59 [–0.69 to – 0.49] mm/h).

For the RNA interference pathway only *qde-2* showed a different reaction norm; other mutant strains did not show any effects.

### Effects of histone demethylation and acetylation

The viable histone acetyltransferase mutant *ngf-1* grew poorly (Figure S6), indicating the critical role this gene plays in normal cellular function. The other mutant, *elp3*, presented reaction norms with slightly lower elevations in all trials. We observed that the *elp3* mutant grew slower at high temperatures. We also observed that its pH optimum dropped to 4.9 from 5.6.

The two putative histone demethylases (*lid2* and *aof2*) had lower elevation in their reaction norms for all environments (Figure S6). However, the reaction norm shape of *lid2* changed as temperature decreased to an optimum of +32.0 °C, but this result could not be validated as there was no significant difference in the change in growth rate between control and *lid2* when temperature increased from +30 to +35 °C. In contrast, *aof2* had a growth rate that was indistinguishable from the control at +35 °C but its growth rate dropped dramatically when temperature increased to +40 °C (Figure S6, Figure 3). The difference to the control in terms of growth rate for *aof2* in +40 °C was –1.26 (–1.85 to – 0.66) mm/h, a drop of 38 %. We validated this result and observed that *aof2* grew slower than the control in the validation experiment at +40 °C as well; a difference of –0.38 (–0.57 to – 0.18) mm/h. However, this growth response was less obvious in the validation experiment where the change in growth rate was not significantly different from the control. Therefore, we conclude that *aof2* and *lid2* are not important to the temperature-dependent responses of *N. crassa*.

Effects of histone acetylation on reaction norms were mainly on reaction norm elevation and only presented slight changes to shape, while histone demethylation affected mainly the elevation of reaction norms.

## Discussion

While epigenetic mechanisms have been shown to be important in certain plastic responses (Slepecky and Starmer, 2009; Bossdorf *et al*., 2010; Herrera *et al*., 2012; Baerwald *et al*., 2015), the extent to which they contribute to phenotypic plasticity or how they maintain homeostasis in organisms facing changing environments has been largely unexplored. By exposing a set of deletion mutants of the filamentous fungus *Neurospora crassa* to a spectrum of controlled environmental parameters, we showed that certain epigenetic modifications have strong effects on plasticity while others do not. In our experiment, epigenetic modifications affected the sensitivity to environmental change and, to a lesser extent, growth of the mutant strain. Modification types did not have a consistent pattern in their effects on phenotype. However, it may be that our classification of epigenetic mechanism was too coarse and this may be why we did not observe a consistent effect. Instead, phenotypic effects were specific to the epigenetic modification in a given environment.

Epigenetic mechanisms clearly played a role in the phenotypic plasticity of growth according to several environmental variables, corroborating recent suggestions concerning the epigenetic control of phenotypic plasticity (Schlichting and Wund, 2014). One of the main findings of this study is that epigenetic modifications were more important for plasticity than average growth rate, i.e., genotypes differed less in terms of their average growth than their variances across the environments. This indicates that plasticities in different environments are caused by different epigenetic mechanisms. There are no uniform plasticity loci, gene expression patterns are context dependent Windig *et al*. (2004), and different epigenetic mechanisms had different effects.

### Histone modifications

Our results suggest that histone modifications play an important role in how *N. crassa* responds to environmental perturbation. Histone modifications H3K36me and H3K4me3 are important in plastic responses to temperature and pH, respectively. *Set-2* is responsible for H3K36 methylation, and the strain lacking a functional form of this gene suffered some developmental deficiencies, i.e., female sterility and production of few conidia (Adhvaryu *et al*., 2005). It also grew slower than the wild type in most environments, but especially so at high temperatures. The optimum growth rate of *set-2* is at +25 °C, while the wild type has an optimum at +35 °C. This indicates that H3K36 methylation is required for the correct expression of genes required at temperatures above +25 °C. In other organisms H3K36 methylation has been associated with transcriptional elongation (Morris *et al*., 2005; Hampsey and Reinberg, 2003); H3K36me is present in the active regions of eukaryotic genomes and its function seems to keep the chromatin of actively-transcribed genes open (Venkatesh *et al*., 2012). It may be that genes expressed under certain environmental circumstances need to be kept in open conformation by H3K36me, and the non-functional strain clearly had a problem at high temperatures. The gene *set-1* responsible for H3K4 trimethylation was important for the response to pH. Previously it has been reported that H3K4me3 is needed for the correct expression of the circadian clock gene *frq* in *N. crassa* (Raduwan *et al*., 2013). In general, H3K4me3 is associated with the 5’-regions of actively-transcribed genes (Pokholok *et al*., 2005; Ardehali *et al*., 2011). In *N. crassa*, *set-1* has a growth phenotype suggesting that H3K4me3 is needed for normal cellular metabolism as well as a specific response to acidic pH. In contrast to *set-1* and *set-2*, the *set-7* mutant strain did not present any phenotypic effect. *set-7* is responsible for H3K27 trimethylation and genes marked with H3K27me3 are silent in *N. crassa* (Jamieson *et al*., 2013). Genes marked with H3K27me3 tend to be less conserved, suggesting that they are only needed in certain environmental conditions. Therefore, it is surprising to observe that the *set-7* mutant strain performed as well as the wild type in our trials. It may be that a lack of repression by H3K27me3 (which allows genes to be expressed) does not prevent a plastic response. The *npf* mutant also lacks H3K27me3 (Jamieson *et al*., 2013), but as it has a very different phenotype than *set-7* and severe growth defects in its phenotype cannot be attributed to lack of H3K27me3. Furthermore, H3K9 methylation seems essential for normal cellular function as *dim-5* mutant lacking H3K9me (Tamaru and Selker, 2001; Tamaru *et al*., 2003) had a severe growth defect in all trials. This phenotype is possibly due to the role of H3K9me in genome integrity (Lewis *et al*., 2010a).

### DNA methylation

DNA methylation in *Neurospora* is directed at regions where histone 3 lysine 9 methylation (H3K9me) is present. H3K9me is required for DNA methylation as *dim-5* lacks both H3K9me and the ability to perform DNA methylation (Tamaru and Selker, 2001; Tamaru *et al*., 2003). DNA methylation can cause gene silencing in *Neurospora* as growth defects of *dmm* mutants were alleviated after the removal of DNA methylation (Honda *et al*., 2010) and DNA methylation can also cause silencing of antibiotic resistance genes (Lewis *et al*., 2010b) but is not required for all gene silencing (Honda *et al*., 2012). However, a complete lack of DNA methylation in the *dim-2* mutant was associated with only a slight response in the 30 % sucrose environment. This is in stark contrast to land plants and vertebrates where DNA methylation appears to be indispensable (Li *et al*., 1992; Rai *et al*., 2006; Xiao *et al*., 2006; Yaari *et al*., 2015). We observed lower elevation of the reaction norms for *dmm-1* and *dmm-2* mutants in all environments and a change in reaction norm shape for *dmm-2* in response to pH (i.e., poor growth at pH 4) suggesting that genes required for this response are silenced as DNA methylation spreads from heterochromatic regions in these mutant strains (Honda *et al*., 2010).

### Histone deacetylation

We examined two different classes of histone deacetylase genes, the Class I histone deacetylases and NAD^+^ dependent Class III histone deacetylases. Class I include the hda genes: *hda-1*, *hda-2*, and *hda-4*. It has been reported that histone H2B is the main target of *hda-1* but it can also deacetylate H3 and is also involved in control DNA methylation (Smith *et al*., 2010; Honda *et al*., 2012), *hda-4* broadly increases acetylation of histones H3 and H4, and no marked effects were reported for *hda-2* (Smith *et al*., 2010). Therefore, it is striking that the phenotypes of *hda-1* and *hda-2* are very similar. These knockout strains presented reaction norms with a similar shape to the wild type in all environmental trials other than osmotic stress (where they grew faster), but with a lower elevation indicating that normal cellular functioning is impaired. Enhanced growth in osmotic stress is surprising, and one interpretation is that there is some cost associated with expressing these genes in this environment. On the other hand, *hda-4* was no different to the control, indicating that it is not involved in phenotypic plasticity in the environments tested here. Class III histone deacetylases include the *nst* genes: *nst-1*, *nst-2*, *nst-4*, *nst-6*, and *nst-7*, these are homologous to the SIR2 family of histone deacetylases (Blander and Guarante, 2004). In *N. crassa*, it has been shown that *nst-1* and *nst-2* are involved in telomeric silencing (4,6, and 7 were not examined) and that *nst-1* deacetylates H4K16 (Smith and Kruglyak, 2008). In other eukaryotes, sirtuin proteins can have other targets than histones (Blander and Guarante, 2004), so for *nst-4*, *nst-6*, and *nst-7* we cannot be certain of their functions. In trials, *nst-1*, *nst-2* and *nst-4* did not present any growth effect in the environments tested. However, the reaction norms of *nst-6* and *nst-7* had pronounced shape changes in sucrose concentration and pH trials. Moreover, the reaction norms of these two mutants were very similar suggesting that they may work in a similar way.

### RNA interference pathway

RNA interference in *Neurospora* may not strictly be an epigenetic mechanism. It is not required for DNA methylation (Freitag *et al*., 2004) and it is not known if RNA-directed epigenetic modifications such as the plant RNA-directed methylation pathway(Matzke and Mosher, 2014) exist or if RNA molecules can mediate epigenetic inheritance like in animals (Rassoulzadegan *et al*., 2006; Rechavi *et al*., 2011; Ashe *et al*., 2012) or other fungi (Qutob *et al*., 2013; Calo *et al*., 2014). Nevertheless, the possibility of RNA-mediated epigenetic effects exists so we included appropriate genes in our examination of the system. We observed that the *N. crassa* Argonaute (Meister, 2013) homolog *QDE-2* was involved in multiple responses to environmental stress, while the two Dicer protein homologs (DCL-1 and DCL-2), *QDE-1*, and *QIP* were not. In *N. crassa*, there are several pathways that generate different kinds of small RNAs; qiRNAs are Dicer-and *QDE-1*-dependent and are involved in the DNA damage response (Lee *et al*., 2009), while the biogenesis of some microRNA-like RNAs (milRNAs) requires *QDE-2* (Lee *et al*., 2010). Other milRNAs are Dicer-dependent and *QDE-2*-independent, some require *QIP* but others do not (Lee *et al*., 2010). While it remains possible that some Dicer-dependent milRNAs are produced by *dcl-1* and *dcl-2* (as there may be some redundancy between these genes: (Catalanotto *et al*., 2004)), our results suggest that those milRNAs whose biogenesis is *QDE-2*-dependent are primarily involved in plastic responses to the environment.

### Histone demethylation and acetylation

Of the remaining genes, *NGF-1* and *ELP3* are believed to be histone acetyl transferases based on their similarity to those genes in yeast (Wittschieben *et al*., 1999; Brenna *et al*., 2012). In *N. crassa*, Brenna *et al*. (2012) showed that NGF-1 is involved in transducing environmental signals, but we found that the *ngf-1* mutant grew very slowly in all environments indicating that key cellular processes are impaired. The *elp3* mutant grows slower and its reaction norms have generally lower elevation, but there was no indication of a shape change. Previously it has been reported for yeast elp3 that it has a temperature-sensitive phenotype (Wittschieben *et al*., 1999). Indeed, we observed that differences in growth between *elp3* and the wild type were largest at +40 °C. However, *elp3* still has the same temperature optimum (+35 °C) as the wild type, suggesting that rather than a temperature response itself, a more fundamental biological process is impaired in the *elp3* mutant strain. The genes *AOF2* and *LID2* are inferred to be histone demethylases. In fission yeast, the *N. crassa* AOF2 protein homolog LSD1 acts as a histone demethylase that demethylates H3K4me and H3K9me (Lan *et al*., 2007). We observed *aof2* reaction norms with lower elevation in salt stress, sucrose concentration, and pH trials. This suggests that certain cellular processes are not functioning normally. The yeast homolog of LID2 also acts as a H3K4 demethylase and interacts with the H3K9 methylation complex (Li *et al*., 2008). The phenotype of the *lid2* mutant is similar to *aof2* in that it had lowered reaction norm elevation in all trials but its reaction norm shape appears similar to wild type.

### Conclusions

In terms of the different environmental parameters tested, we observed that epigenetic mechanisms in *N. crassa* play a much greater role in the response to temperature and pH changes than they do to shifts in sucrose concentration and osmotic stress. This can be explained by the ecology *Neurospora*, and realizing that *N. crassa* is a saprotrophic fungus that is found in dead plant matter (Jacobson *et al*., 2006) or as an endophyte under certain conditions (Kuo *et al*., 2014). Temperature changes are the most common environmental variable that organisms experience and pH changes are likely to occur as the fungus encounters different substrates in nature. It may be that *N. crassa* rarely encounters elevated NaCl levels in a terrestrial environment and has not evolved a plastic response to it. In the sucrose concentration trial we examined how the level of available nutrients and osmotic stress affect the growth of *N. crassa*, and it would be interesting to investigate how the fungus responds to different types of carbon sources and whether those responses are under epigenetic control.

Another question that requires investigation is whether the plastic responses we have detected are heritable. It has been observed that maternal or transgenerational effects can be mediated mechanistically by epigenetic changes. In plants, DNA methylation and RNA-directed methylation in particular have been implicated in transgenerational inheritance (Luna *et al*., 2012; Luna and Ton, 2012). In fruit flies, histone modifications have also been linked to transgenerational inheritance where H3K27 and H3K9 methylation regulate offspring lipid content in response to paternal diet (Öst *et al*., 2014).

Our results show that that epigenetic mechanisms are involved in plastic responses of *N. crassa* and that histone methylation is likely to be main mechanism along with small RNAs that are dependent on *QDE-2*. We suggest that epigenetic mechanisms are likely to be important mediators of plastic responses. Epigenetic mechanisms may also facilitate evolutionary adaptation via phenotypic plasticity, as suggested by models (Lande, 2009; Chevin *et al*., 2010; Draghi and Whitlock, 2012) and experiments (Schaum and Collins, 2014; Lind *et al*., 2015). The exact effects will depend on whether plasticity occurs within or across generations, which will be an important topic of research in the future.

## Acknowledgments

We acknowledge the Fungal Genetics Stock Center (Kansas City, Missouri USA) for providing the *Neurospora* strains and Mr. Juha Ahonen for constructing the race tube racks. We would like to thank Eric Selker and Jouni Laakso for comments on the manuscript. Michael Hardman of Lucidia checked the English. This research is supported by the Academy of Finland grant no. 274769 to IK and no. 278751 to TK and the Centre of Excellence in Biological Interactions of University of Jyväskylä.

---

**Figure S1:**
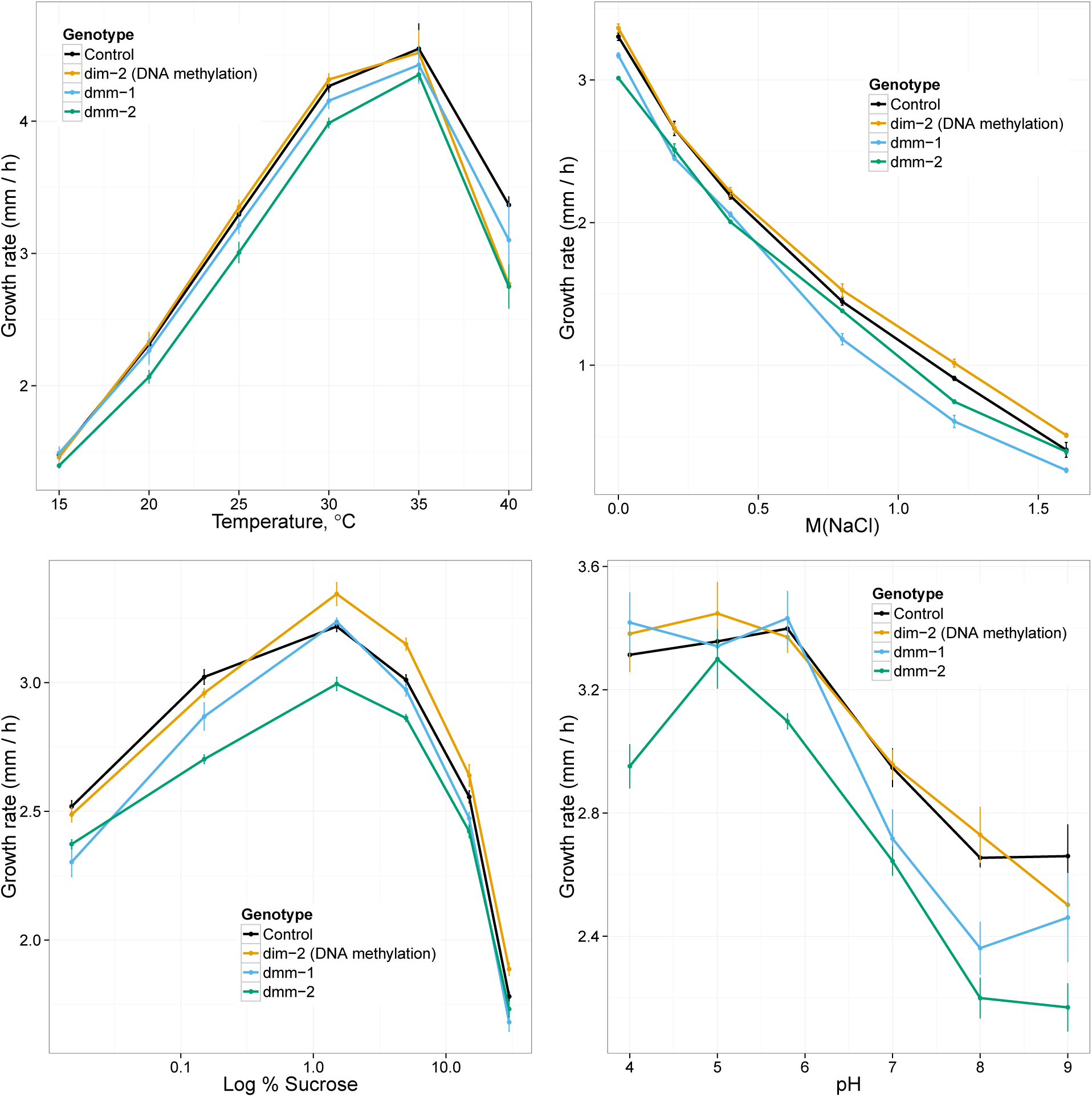
Reaction norms for DNA methylation mutants in the four environmental parameters. Reaction norms are colored according to mutant strain and the control as indicated in the legend. Error bars are ± SE.

**Figure S2:**
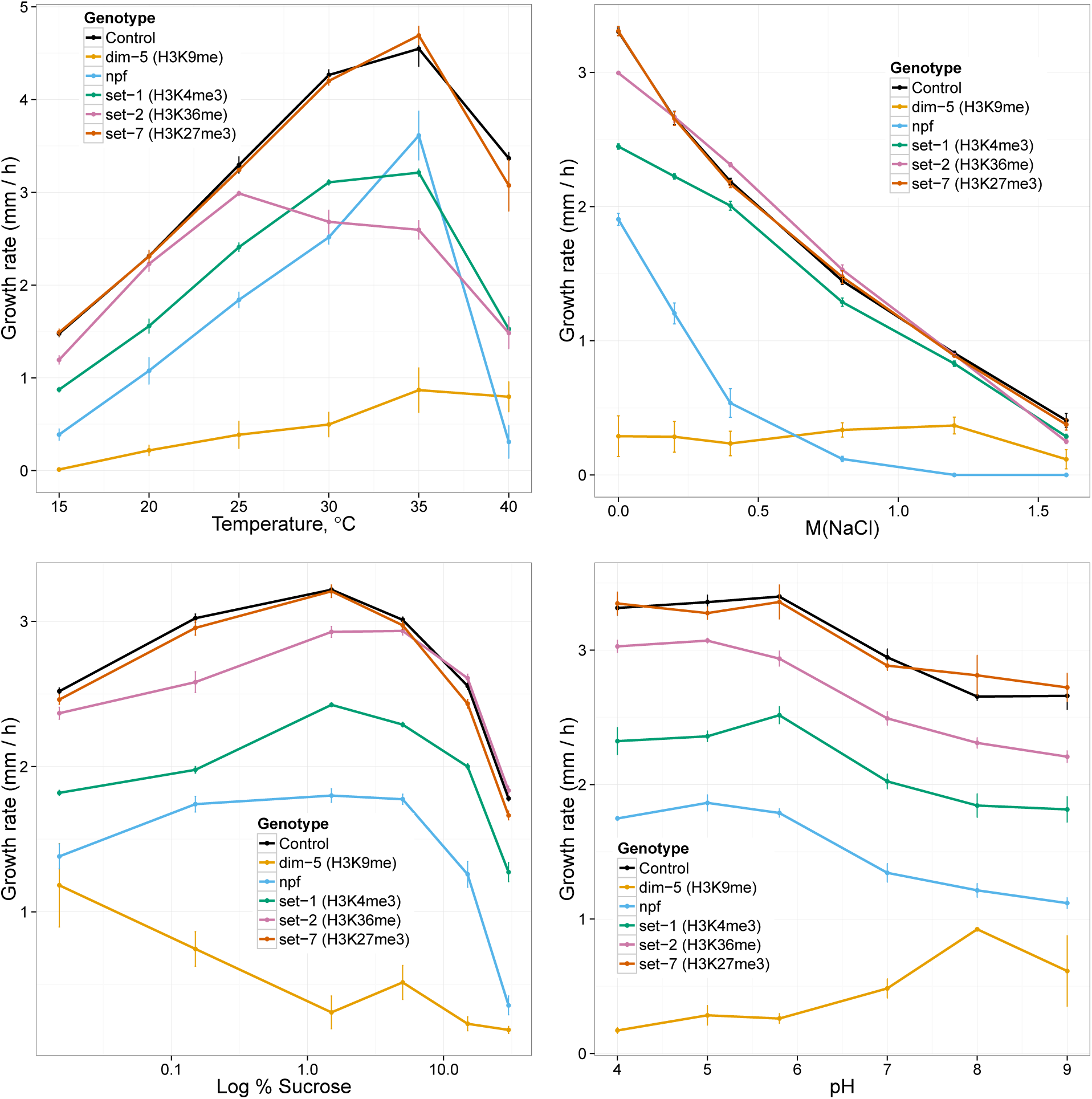
Reaction norms for histone methylation mutants in the four different environmental parameters. Reaction norms are colored according to mutant strain and the control as indicated in the legend. Error bars are ± SE.

**Figure S3:**
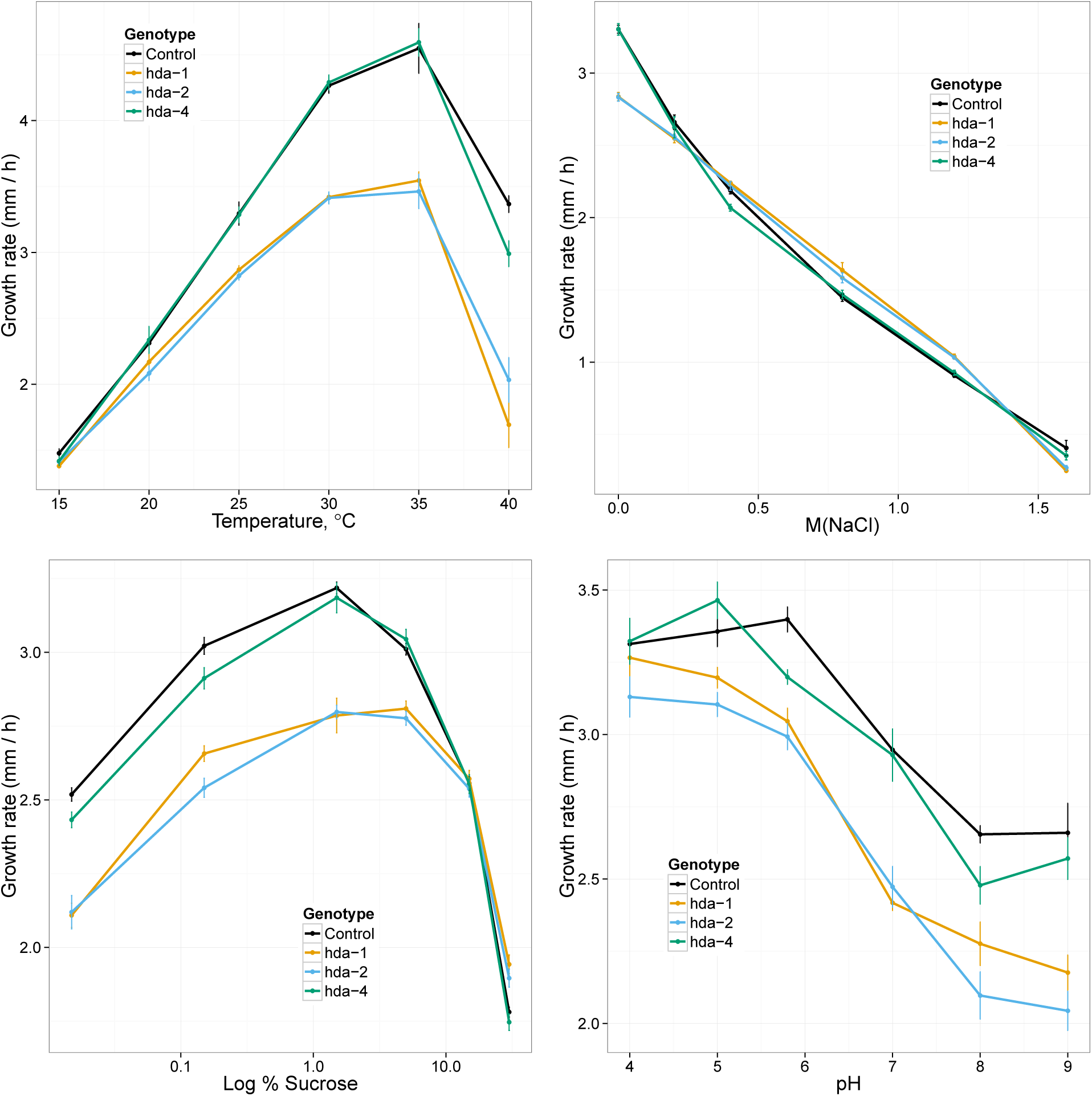
Reaction norms for histone deacetylation mutants in the four environmental parameters. Reaction norms are colored according to mutant strain and the control as indicated in the legend. Error bars are ± SE.

**Figure S4:**
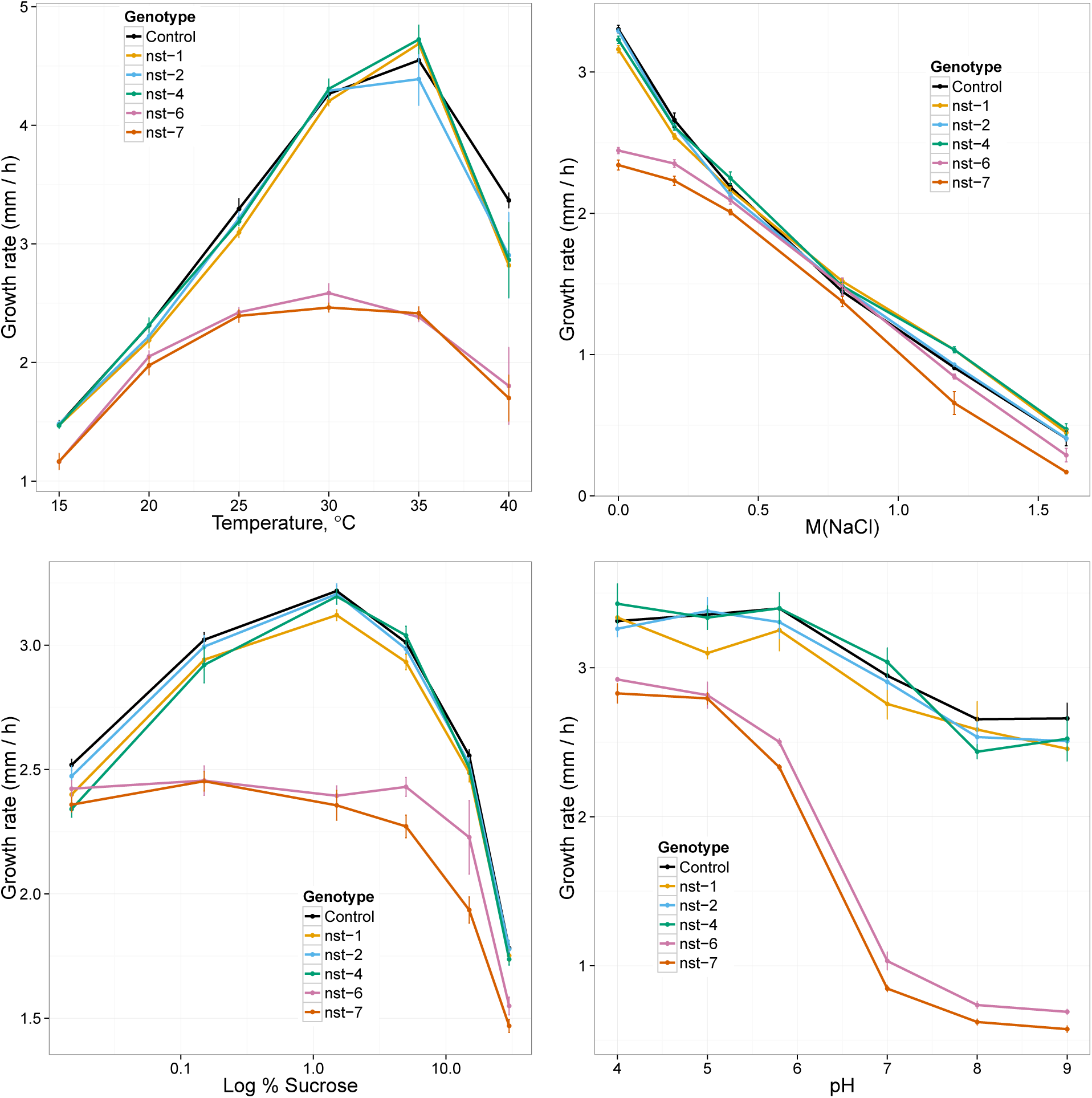
Reaction norms for histone deacetylation, type III mutants in the four different environmental parameters. Reaction norms are colored according to mutant strain and the control as indicated in the legend. Error bars are ± SE.

**Figure S5:**
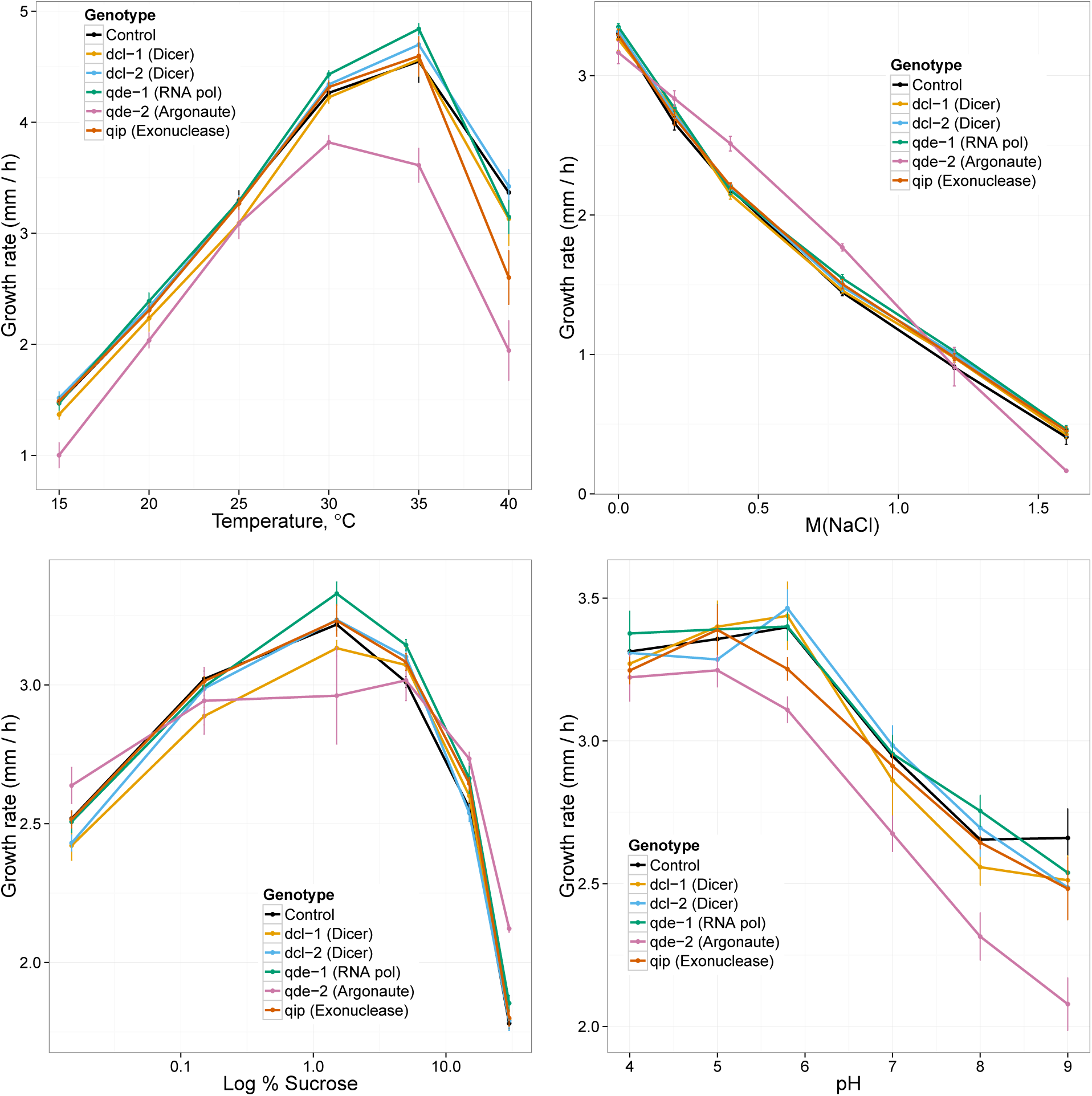
Reaction norms for RNA interference mutants in the four different environmental parameters. Reaction norms are colored according to mutant strain and the control as indicated in the legend. Error bars are ± SE.

**Figure S6:**
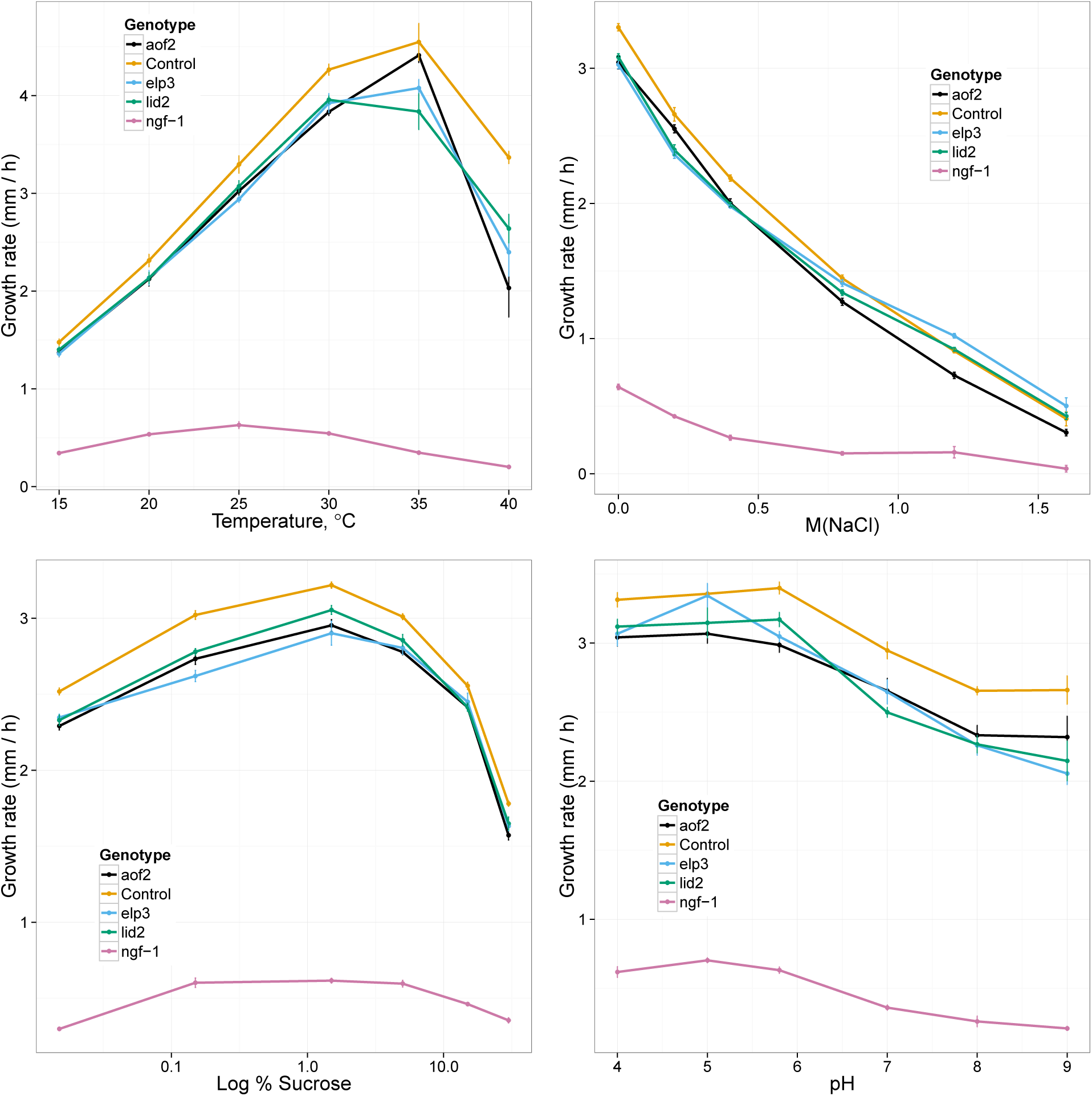
Reaction norms for histone acetylation and putative histone demethylase mutants in the four different environmental parameters. Reaction norms are colored according to mutant strain and the control as indicated in the legend. Error bars are ± SE.

**Table S1:** *Neurospora* strains used in this study. Strains used in the reaction norm experiment are indicated as are the strains used in the validation experiments. Strains marked as preparation were used for generating some of the strains used in the experiments.

| Genotype | Mating type | Source | Strain ID | Experiment |
| --- | --- | --- | --- | --- |
| $\Delta dim-2::hph$ | *mat a* | Selker lab | N1850 | reaction norms |
| $\Delta dmm-1::hph$ | *mat a* | FGSC | 14504 | reaction norms |
| $\Delta dmm-2::hph$ | *mat a* | FGSC | 11100 | reaction norms |
| $\Delta dim-5::hph$ | *mat a* | this study | K1 | reaction norms |
| $\Delta set-1::hph$ | *mat a* | this study | K2 | reaction norms |
| $\Delta set-2::hph$ | *mat a* | FGSC | 15504 | reaction norms |
| $\Delta set-7::hph$ | *mat a* | FGSC | 11182 | reaction norms |
| $\Delta npf::hph$ | *mat a* | FGSC | 13915 | reaction norms |
| $\Delta nst-1::hph$ | *mat a* | FGSC | 12403 | reaction norms |
| $\Delta nst-2::hph$ | *mat a* | FGSC | 12078 | reaction norms |
| $\Delta nst-4::hph$ | *mat a* | FGSC | 11165 | reaction norms |
| $\Delta nst-6::hph$ | *mat a* | FGSC | 22669 | reaction norms |
| $\Delta nst-7::hph$ | *mat a* | FGSC | 16002 | reaction norms |
| $\Delta hda-1::hph$ | *mat a* | FGSC | 12003 | reaction norms |
| $\Delta hda-2::hph$ | *mat a* | FGSC | 11158 | reaction norms |
| $\Delta hda-4::hph$ | *mat a* | FGSC | 11175 | reaction norms |
| $\Delta qde-1::hph$ | *mat a* | FGSC | 11156 | reaction norms |
| $\Delta qde-2$ | *mat a* | Selker lab | N2271 | reaction norms |
| $\Delta dcl-1::hph$ | *mat a* | FGSC | 15892 | reaction norms |
| $\Delta dcl-2::hph$ | *mat a* | FGSC | 11155 | reaction norms |
| $\Delta qip::hph$ | *mat a* | FGSC | 12130 | reaction norms |
| $\Delta aof2::hph$ | *mat a* | FGSC | 11964 | reaction norms |
| $\Delta elp3::hph$ | *mat a* | FGSC | 11976 | reaction norms |
| $\Delta lid2::hph$ | *mat a* | FGSC | 11876 | reaction norms |
| $\Delta ngf-1::hph$ | *mat a* | FGSC | 16229 | reaction norms |
| *wt* | *mat a* | FGSC | 4200 | reaction norms |
| *wt* | *mat A* | FGSC | 2489 | validation |
| $\Delta dim-2::hph$ BC<sub>5</sub> 2489 | *mat A* | this study | K3 | validation |
| $\Delta dmm-2::hph$ BC<sub>5</sub> 2489 | *mat A* | this study | K4 | validation |
| $\Delta hda-1::hph$ BC<sub>5</sub> 2489 | *mat A* | this study | K5 | validation |
| $\Delta qde-2$ BC<sub>5</sub> 2489 | *mat A* | this study | K7 | validation |
| $\Delta qip::hph$ BC<sub>5</sub> 2489 | *mat A* | this study | K8 | validation |
| $\Delta aof2::hph$ BC<sub>5</sub> 2489 | *mat A* | this study | K9 | validation |
| $\Delta lid2::hph$ BC<sub>5</sub> 2489 | *mat A* | this study | K10 | validation |
| $\Delta set-7::hph$ BC<sub>6</sub> 2489 | *mat A* | this study | K11 | validation |
| $\Delta set-1::hph$ | *mat A* | FGSC | 15828 | preparation |
| $\Delta dim-5::hph$ HET | *mat a* | FGSC | 15885 | preparation |

**Table S2:**
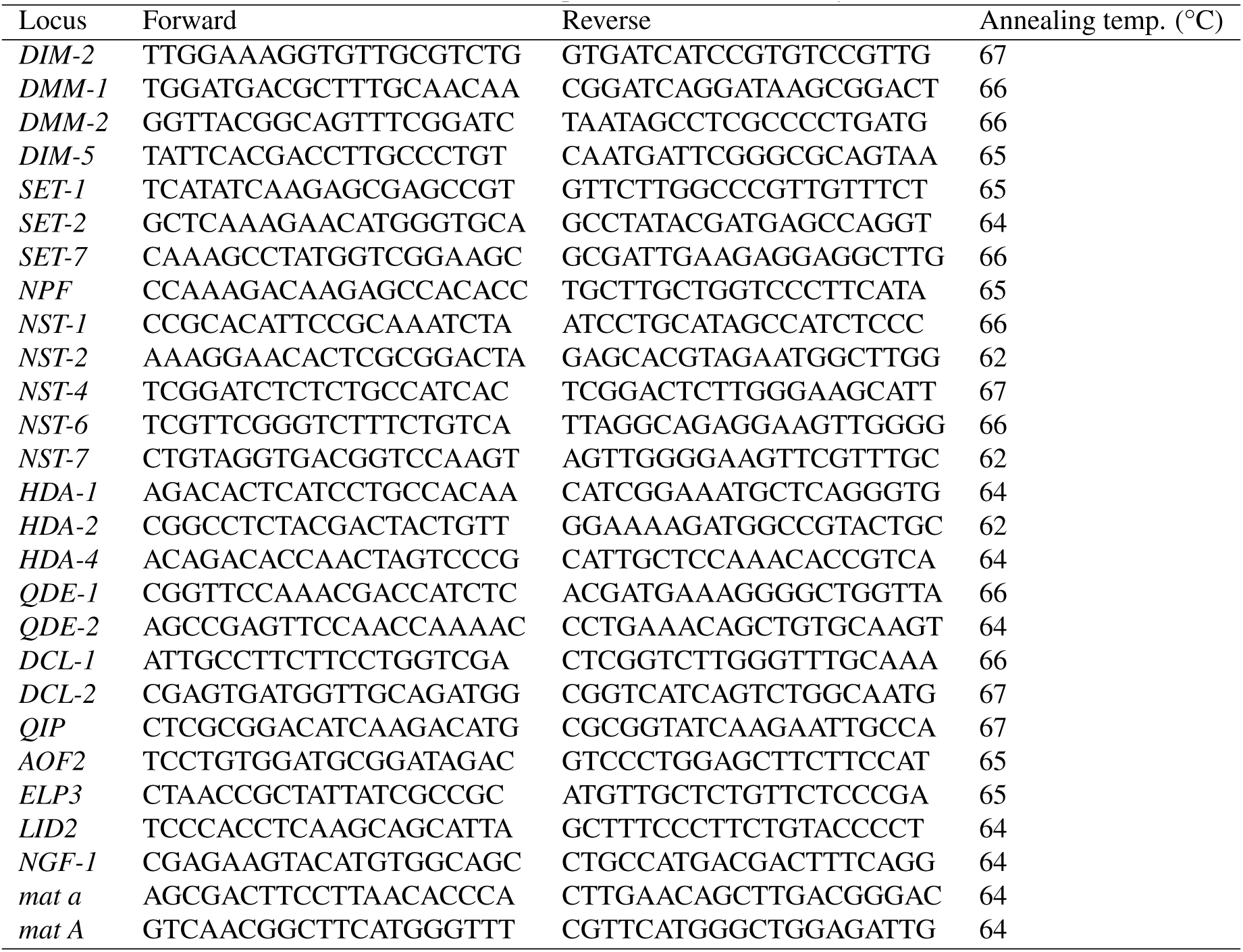
PCR primers used in this study.

